# Daratumumab induces mechanisms of immune activation through CD38+ NK cell targeting

**DOI:** 10.1101/849265

**Authors:** Domenico Viola, Ada Dona, Enrico Caserta, Estelle Troadec, Emine Gulsen Gunes, Francesca Besi, Tinisha McDonald, Lucy Ghoda, James F Sanchez, Jihane Khalife, Marianna Martella, Chatchada Karanes, Myo Htut, Xiuli Wang, Michael Rosenzweig, Arnab Chowdhury, Douglas Sborov, Rodney R Miles, Paul J. Yazaki, Stephen J Forman, John Shively, Guido Marcucci, Steven T Rosen, Jonathan J Keats, Amrita Krishnan, Flavia Pichiorri

**Author notes:** These authors equally contributed to this work (Domenico Viola, Ada Dona). **Corresponding Authors:** Flavia Pichiorri, Department of Hematologic Malignancies Translational Science, City of Hope, 1500 E. Duarte Rd., Duarte, CA 91010; Amrita Krishnan, Department of Hematology and Hematopoietic Cell Transplantation, City of Hope, Duarte, CA 91010.

## Abstract

Daratumumab (Dara), a multiple myeloma (MM) therapy, is an antibody against the surface receptor CD38, which is expressed not only on plasma cells but also on NK cells and monocytes. Correlative data have highlighted the immune-modulatory role of Dara, despite the paradoxical observation that Dara regimens decrease the frequency of total NK cells. Here we show that, despite this reduction, NK cells play a pivotal role in Dara anti-MM activity. CD38 on NK cells is essential for Dara-induced immune modulation, and its expression is restricted to NK cells with effector function. We also show that Dara induces rapid CD38 protein degradation associated with NK cell activation, leaving an activated CD38-negative NK cell population. CD38+ NK cell targeting by Dara also promotes monocyte activation, inducing an increase in T cell costimulatory molecules (CD86/80) and enhancing anti-MM phagocytosis activity ex-vivo and in vivo. In support of Dara’s immunomodulating role, we show that MM patients that discontinued Dara therapy because of progression maintain targetable unmutated surface CD38 expression on their MM cells, but retain effector cells with impaired cellular immune function. In summary, we report that CD38+ NK cells may be an unexplored therapeutic target for priming the immune system of MM patients.

## Introduction

Daratumumab (Dara) is a humanized IgG1 (ĸ subclass) antibody against the highly expressed plasma cell (PC) receptor CD38.^1–5^ It has been approved by the Food and Drug Administration for the treatment of relapsed and newly diagnosed multiple myeloma (MM).^1, 3–8^ The main anti-MM effect of Dara has thus far been attributed to its ability to target the MM cells by inducing immune activation cell killing,^9^ but unfortunately, despite its significant efficacy, relapse or resistance remains an issue. Survival following relapse from a Dara regimen is particularly poor and ranges from 9-12 months.^10^ In support of its function as an ectoenzyme, we and others recently reported that a fraction of the entire CD38 molecule is actively internalized when cancer cells, including MM cells, are treated with CD38-specific antibodies.^11, 12^ The correlation between CD38 surface levels on the MM cells and response to Dara treatment remains controversial.^13, 14^ Whereas some researchers reported a significant downregulation of CD38 expression on the surface of the MM-PCs in patients progressing under Dara treatment,^13^ others have instead shown that detection of CD38 on these cells was hindered by competitive binding of Dara, which resulted in a non-specific MM CD38-negative population.^15^ A restoration of CD38 expression on the cancer cells six months after Dara discontinuation has been also described.^13^ Despite the importance of CD38 expression on the myeloma cells, correlative studies have highlighted that MM patients who participated in Dara monotherapy trials (SIRIUS and GEN501) show significant lower levels of total NK cells but an increase in a CD38(-), activated NK cell population (CD69+), associated with an increase in CD8+ T-cell activation after two months of treatment.^16^ Although a recent published study has highlighted the possible effect of Dara in killing CD38+ NK cells with subsequent expansion of a more active CD38(-) NK cell population,^17^ the expansion of this population have not been observed in the clinic, and CD38 signaling has mainly been implicated in mechanisms of NK cell and in general Th1 activation signaling.^18–21^ By using primary patient samples in conjunction with ex-vivo and in vivo testing, we report that Dara binding to CD38 in NK cells induces its internalization and concomitant activation of a CD38+ NK cell population, a first step which we believe is essential in inducing a subsequent immune activation cascade against cancer cells. We also report that Dara progressing patients retain targetable CD38 surface expression on their cancer cells but impaired Dara induced immune killing ability on their effector cells.

## Results

### Dara-induced MM cell killing through CD38+/CD16+NK cells

Confirming previously published data with single-agent Dara,^16^ our data show that refractory patients responding to Dara combinatorial therapy (actively in treatment) display a significantly lower total NK cell frequency in their peripheral blood compared to that in Dara-naïve refractory patients (RRMM) (Fig.1A). We then decided to investigate whether, despite the reduction of the frequency of this population, these cells still play a role in Dara anti-MM activity. The effect of NK cells was tested in NSG mice engrafted with CD38+ MM cells (MM.1S Gfp/Luc+). Specifically, mice were injected with 5×10^6^ cells, and two weeks after injection, mice with comparative tumor burden were randomly separated in four different treatment groups. Six mice were used for each treatment group and co-injected with the following: 1×10^6^ healthy donor-derived peripheral blood mononuclear cells (PBMCs) plus Dara (gr1) or a non-MM specific humanized IgG1 (ĸ subclass) control antibody trastuzumab (C-IgG) (gr2); or PBMCs depleted of the total NK population plus Dara (gr3); or C-IgG (gr4) (Fig.1B). Each treatment was repeated once a week for three weeks. Our data show that mice treated with PBMCs+Dara have a significantly longer survival compared to that in mice treated with PBMCs+C-IgG (p=0.001), and that depletion of the NK cell population resulted in shorter survival of all the mice injected with PBMCs depleted of total NK cells (PBMCs without NK cells) and treated with Dara, compared to the mice injected with total PBMCs+Dara (p=0.015) (Fig.1C).

**Fig. 1.**
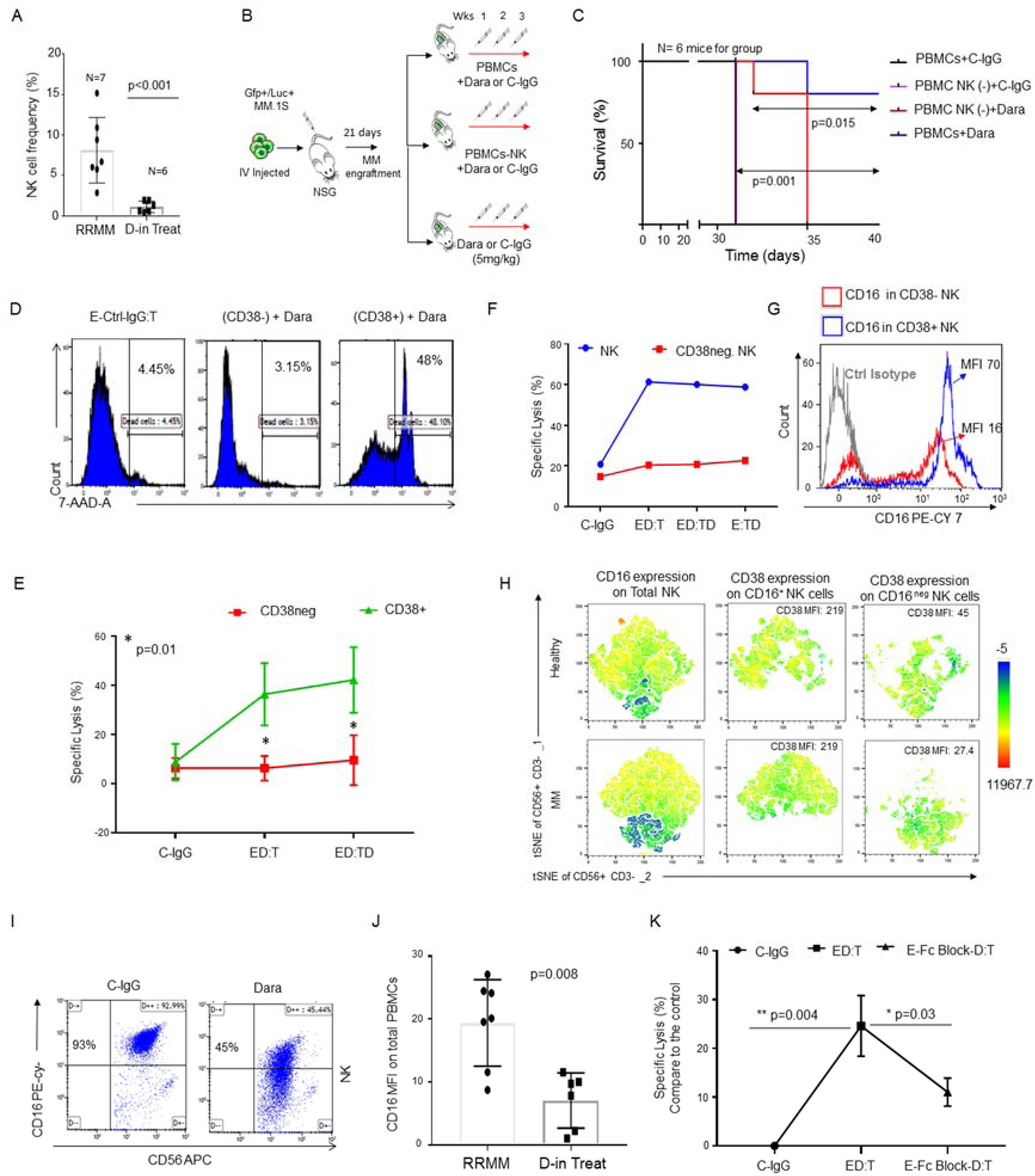
CD38+ NK cells are essential for Dara-induced killing of MM cells. **A)** Bar graphs showing NK cell reduction in the PBMCs isolated from refractory MM patients (RRMM) and refractory MM patients actively on Dara treatment (D-in Treat) (p<0.001). Mann-Whitney-Wilcoxon test was performed; **B)** Schematic illustration of the animal treatment schedule. 5×10^6^ MM.1S GFP+/Luc+ cells were injected intravenously (IV) into NSG mice. On day 13, after engraftment reached approximately ≥2×10^6^ photons/sec/cm^2^/sr, mice were randomly distributed into 4 experimental groups and injected three times once a week by intravenous injection with PBMCs or PBMCs depleted of total NK cells, each isolated from healthy donors (HDs), and co-injected with Dara or trastuzumab (C-IgG) (5 mg/kg); **C)** Survival curves showing shorter survival of mice where human PBMCs were depleted of the NK cell fraction and treated with Dara (Dara PBMCs-NK cells), compared to Dara-treated total PBMCs (p=0.015); **D)** Representative killing assay of PBMC CD38+ and CD38(-) fractions isolated from HD PBMCs (Effector, E) treated for 24hr with C-IgG or Dara (ED) (10 or 100 µg/ml), washed, then co-cultured with Gfp+ MM.1S (Target, T) at a ratio of 8:1 for 12hr; **E)** Line graph reporting the percentage of dead cells as 7-AAD-positive among target cells (MM.1S GFP+ cells) of at least n=3 HDs repeated in triplicate; **F)** Killing assay in terms of specific lysis (%) by total NK cells or CD38(-) NK cells (E) isolated from HDs and incubated overnight with C-IgG and Dara, washed, and co-cultured with Gfp+ MM.1S cells (T) for 4hrs; **G)** Overlay histogram showing that CD16 levels are almost 80% less on the surface of CD38(-) NK cells compared to that in the CD38+ NK cell population in the blood of HDs.; **H)** Mass cytometry analysis showing CD38 distribution in the CD16+ NK cell population in the PBMCs isolated from a healthy (n=1) and a patient (n=3) donor (n=4). **I)** Representative flow cytometry of CD16 surface expression of NK cells after an overnight treatment with C-IgG or Dara; **J)** Bar graph showing CD16 levels in non-refractory MM patients actively on Dara treatment (D-in Treat) compared to that in untreated RRMM patients (p=0.008); **K)** Blocking of the FcR on the effector cells before Dara incubation significantly reduced MM cell killing (p=0.03). The experiment was repeated in 4 independent HDs, each repeated in triplicate. Values represent the mean ± SD; p values were calculated using t-test (tails = 2, type = 1).

We then investigated whether CD38 surface expression on immune effectors was essential for Dara-induced cell killing. PBMCs (effectors, E) were pretreated with Dara (ED), washed, and incubated with the target (T) cells (ED:T). Our data show that only CD38+ effector cells (T cells 69.48%, monocytes 9.14%, NK cells 14.13%, B cells 7.25%) induced MM killing upon Dara treatment (Fig.1D,E, Sup.Fig.1A) when pretreated with Dara (ED:T), whereas the PB CD38-negative fraction (PB-CD38(-)), which includes a subset of NK cells (∼1.3%), CD38(-) T cells (∼95.22%), and monocytes (∼3.58%), were unable to induce MM killing under the same conditions (Fig.1D,E). To avoid the possibility that the PB-CD38(-) fraction was inducing less MM killing due to the presence of a less enriched total NK cell population, we performed a killing assay using purified NK cells. Consistent with the importance of CD38 expression on the surface of immune effector cells, the same effect was also observed in purified total NK cells in which we observed significant MM cell killing at different effector-target ratios, even when MM cells were incubated with CD38+NK cells (Sup.Fig.1 B,C) (Fig.1F). Conversely, CD38(-) NK cells did not induce killing in the same experimental conditions (Fig.1F). Since CD16 expression is critical for NK cell activation against cancer cells by antibody dependent cellular cytotoxicity (ADCC), we investigated whether surface CD16 levels may differ in CD38+ and CD38(-) NK cell populations. Our data show that CD38(-) NK cells have almost 80% less surface CD16 surface expression (MFI 16) compared to that on CD38+NK cells (MFI 70), as analyzed in 4 independent donors (p<0.0001) (Fig.1G). Mass cytometry analysis also confirmed that in both healthy and patient donors, CD38 distribution in CD56+NK cells is mainly restricted to the CD16+ population (Fig.1H). We then assessed whether Dara could directly engage CD16 on the surface of NK cells even in absence of CD38+ target cells. We observed surface CD16 down-modulation when NK cells were treated overnight with Dara (Fig.1I), and the same result was observed in cells obtained from patients actively under Dara treatment (Fig.1J). Interestingly, when the Dara-treated NK cells were washed twice to eliminate the excess of antibody and incubated with the target (CD38+ MM cells, ED:T), an antibody binding the same epitope, CD38-Mono-PE, could not detect CD38 on the surface of MM cells, in contrast to detection following control IgG (C-IgG) but similarly to when MM cells were directly exposed to Dara (ED:TD) (Sup. Fig.1D), clearly showing that when the Fc of Dara is first bound to CD16, the Fab of Dara can efficiently recognize CD38 on the surface of MM cells. Considering the influence of the Fc receptors in bringing cancer cells and immune cells in proximity for activating the immune effector function,^22, 23^ we investigated whether Dara-induced MM cell killing could be hindered by using a total FcR blocker. Blocking the FcR on effector cells (E-FcBlock-D:T) almost completely reverted MM cell killing by Dara using 4 different donors (Fig.1K).

### Dara induces CD38+ NK cell activation

Since CD38+ effector cells appear crucial for the engagement of the Dara-induced anti-MM immune response but previously published data have shown that Dara-treated patients are left with an activated CD69+CD38(-) NK cell population, we decided to explore this apparent discordancy ex-vivo. Accordingly, we observed CD69 to be considerably upregulated in total NK cells when PBMCs were treated overnight with Dara (Sup.Fig.2A). CD69 upregulation in NK cells was also observed in the CD38+ PBMC fraction, but not in the PBMC CD38(-) population (Sup.Fig.2A). To assess whether Dara could have a direct activating effect on NK cells, we performed the same experiment using pure CD38+ and CD38(-) NK cell populations (Sup.Fig.2B). We observed a strict significant up-regulation in CD69 expression in the CD38+ and in the total NK cell population after overnight treatment with Dara, which was not observed in the CD38(-) fraction (p<0.001) (Fig.2A,B). Because previous data have shown that CD38 and CD16 are functionally dependent to induce mechanisms of NK cell activation^24^ and that Dara could induce fratricide of NK cells by its concomitant binding (bridging) to the Fc receptor and their surface CD38,^17^ we investigated whether targeting CD38 using only the Dara Fab or Fc fragments (Sup.Fig.2C) would still induce NK cell activation. One hour of incubation with the Dara Fab fragment cannot induce NK cell activation (CD69+ cells: 51% from Dara versus 15% from Fab) and degranulation (CD107a+ cells: 24% in Dara versus 0.66% from Fab) (Fig.2C). However, the addition of the purified Dara Fc fragments 30 min after the full saturation of CD38 on NK cells by Fab (Fig. 2D) completely restored the activation observed using the intact antibody (CD69+: Dara 51% versus Fab+Fc 54%; CD107a+: Dara 24% versus Fab+Fc 17%) (Fig. 2E), excluding the possibility of a possible mechanisms of ADCC between NK cells in which both receptors were occupied but not physically connected (Fig.2D). NK cell activation was not observed when Trast, which shares with Dara the same Fc fragment, was used in the same experimental condition, but elevated CD69 and CD107 was again observed when Dara was added (Fig.2E). Western blot analysis revealed that up to 48hrs of treatment with Dara Fab did not induce CD38 down-modulation, but the addition of an intact Dara Fc again decreased CD38 levels comparable to those in Dara-treated NK cells (Fig.2F). Interestingly, we observed CD38 mRNA upregulation in NK cells upon incubation with Dara at 24 hours, and no significant changes were observed at 48hrs of treatment (Fig.2G), supporting that ex-vivo Dara treatment does not select a CD38(-) NK cell population, which in patients retains lower CD38 mRNA levels compared to that in CD38+ NK cells (Sup. Fig.2D). Since previous data have shown that modulation of CD38 signaling pathways induces increases in IFN-γ and GM-CSF levels in NK cells,^24^ we investigated whether Dara could also modulate the release of these cytokines. We found an increase in IFN-γ and GM-CSF production in NK cells treated at different time points (1-16 hrs) at different Dara concentrations (1-100 µg/ml) (Fig.2H,I). As expected, IFN-γ and GM-CSF increases were associated with upregulation of the NK cell degranulation marker (CD107a) (Fig.2J).

**Fig. 2.**
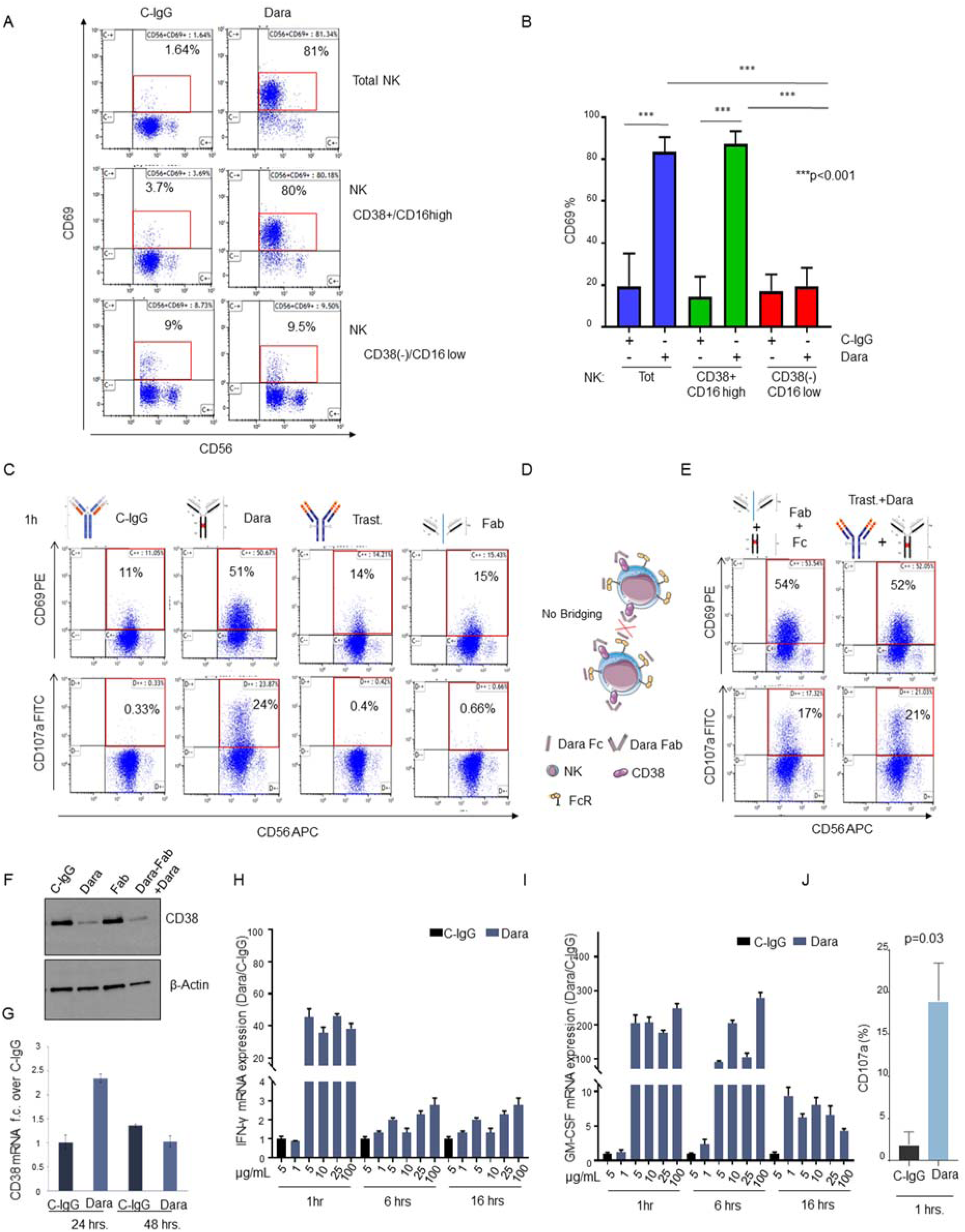
Dara induces direct CD38+ NK cell activation. **A)** Representative flow analysis of CD69 on total NK, CD38+NK and CD38(-) NK cell fractions isolated from a healthy donor upon overnight Dara treatment; **B)** Bar graph representing the median ±SD of CD69 expression in total, CD38(-) and CD38+ NK cells treated with Dara or C-IgG (p<0.001). Each experiment was repeated using three healthy donors in triplicate; **C,E)** Flow cytometry analysis of CD69 (top panels) and CD107a (bottom panels) in total NK cells treated for 1hr with C-IgG, Dara, Dara Fab (Fab), Trast, Dara Fab + Dara Fc (Fab+Fc), or Trast+Dara at 10 µg/mL for 1 hr; **D)** Graphical Illustration showing the fully saturated CD38 on NK cells after the addition of the purified Dara Fc fragment for 30 min, which followed the addition of Dara Fab, excluding a possible bridging between NK cells; **F**) Western Blot analysis showing CD38 protein levels in NK cells treated with either C-IgG (10 μg/mL), Dara (10 μg/mL), Dara-Fab or Dara-Fab+Dara for 48 hrs; **G**) Graph bar showing CD38 mRNA expression levels in primary NK cells treated with C-IgG as detected by q-RT-PCR normalized for GAPDH at 24 and 48 hrs; **H,I)** INF-γ and GM-CSF mRNA expression levels in NK cells under Dara treatment at different concentrations (from 1 µg/ml to 100 µg/ml) and different time points (1-16 hrs) compared to C-IgG; GADPH mRNA was used for normalization; **I)** Graph bar showing surface CD107a expression (%) in NK cells after 1hr of Dara or C-IgG treatment (p=0.03). Results are representative of the average of at least 3 healthy donors.

### Dara induces CD38 protein degradation through the auto-phagosome apparatus

Flow cytometry analysis using anti-CD38-Multi-FITC, a non-competitive antibody, shows that CD38 on the surface of NK cells from healthy donors (HDs) is strongly downregulated upon Dara treatment, from 70% to 40% after 1hr and to 18% after overnight treatment, an effect not observed with C-IgG (Fig.3A). Despite this surface downregulation, the total CD38 protein content upon 1 hour of incubation was unaffected (Fig.3B). Regardless of differences in CD38 levels, a decrease in CD38 protein was observed as soon as 6hrs in all HDs tested (n=13; median age 56), in contrast to levels in C-IgG–treated NK cells (p=0.008) (Fig.3B,C). Since Sullivan et al. showed that in MM patients Dara causes CD38 degradation in red blood cells,^25^ and because we recently published that the Dara/CD38 complex can be internalized in the MM cells,^26^ we investigated whether Dara binding on the CD38+ NK cells could cause Dara/CD38 complex internalization and subsequent degradation. We used our recently published Dara conjugated to Alexa-Fluor 647 (DARA-AF-647).^26^ We first blocked DARA-AF-647 active NK cell internalization using ice and assessed maximum surface signal after 120 min of incubation; minimum surface signal in the same cells was alternatively assessed after surface acid wash (a.w.). NK cells were then incubated with DARA-AF-647 at 37°C at different time points (15–120 minutes), and the internalized fluorescence left after a.w. was compared to the minimum and maximum surface signals (Fig. 3D,E). We observed a rapid DARA-AF-647 internalization in CD38+ NK cells, a signal that reached a maximum internalization level at 60 min and remained elevated at 120 min after incubation. This result is aligned with the stability in total CD38 protein levels in the first few hours of treatment (Fig.3C). Our data show that Dara treatment induces an increase in intracellular Ca^2+^ mobilization in NK cells, with a 40% increase in magnitude after 30 seconds of Dara addition (Fig.3F). Since we previously published that CD38 can be internalized though the auto-phagosome apparatus, we assessed whether CD38 protein down-modulation in NK cells could be linked to lysosomal degradation. Dara treatment induces a decrease of the phagosome marker microtubule-associated protein light chain 3 (LC3), and increase in GLUT4, supporting an increase in glucose uptake induced by autophagy and lysosomal degradation (Fig.3G). We also observed that primary NK cells treated with the autophagy inducer rapamycin (10, 50 µM) for 6 hours in the presence of either Dara or C-IgG show CD38 protein downregulation in a dose-dependent manner, an effect that was further potentiated by the addition of Dara (Fig.3H). We then investigated whether CD38 protein degradation upon Dara binding could be reverted. When NK cells were treated overnight with the autophagy inhibitor wortmannin (Wort), we observed a significant rescue of CD38 protein degradation upon Dara binding compared to that in the NK cells treated only with Dara (Fig.3I,J). Consistent with these data, CD38 surface expression was also partially rescued by the addition of Wort. (Fig.3K).

**Fig. 3.**
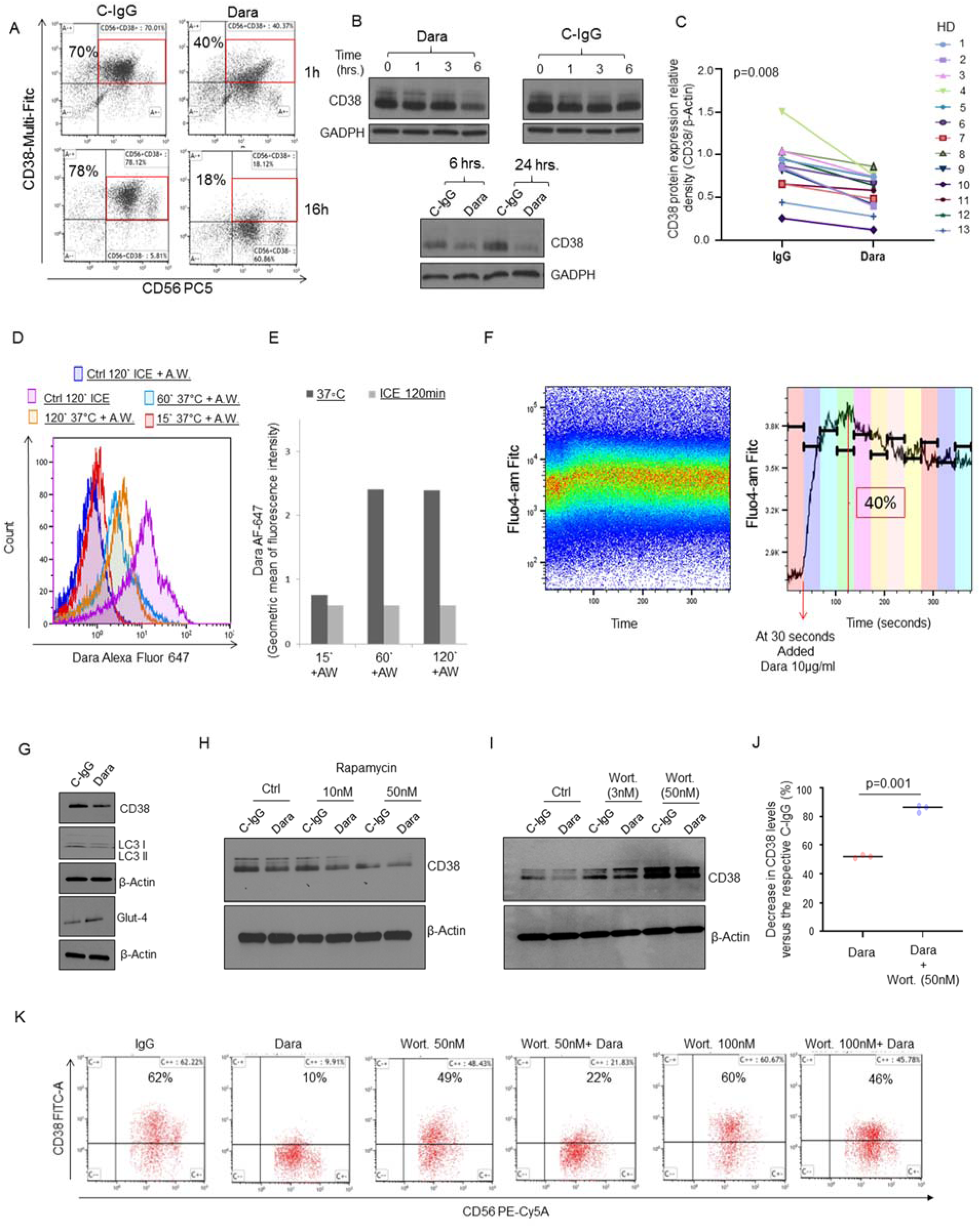
Dara induces CD38 protein reduction in NK cells and Ca^2+^ mobilization. **A)** CD38 surface expression on NK cells at 1hr and 16hrs as detected by flow cytometry, showing down-regulation upon Dara treatment at 1hr and 16hrs compared to C-IgG; **B)** Time course of 0-1-3-6hrs and 6-24hrs as indicated for NK cells isolated from HDs and treated with Dara (10 μg/mL) or IgG (10 μg/mL) and as assessed by Western Blot analysis of CD38 protein level expression. **C)** Thirteen tumor-free donors (HD) were analyzed, and CD38 levels were assessed by densitometry analysis using β-actin as internal housekeeping. The Shapiro-Wilk test suggests that the samples are generated from a Gaussian distribution (a=0.05). Welch t-test statistic = 2.5963 with p=0.008; **D)** Flow cytometry analysis showing increased CD38 internalized in NK cells incubated at 37°C at different time points (15’-60’-120’) with Dara-AF-647 + acid wash (a.w.). Graphical representation of geometric mean of Dara-AF-647 internalized signal in NK cells incubated at different time points (15’ to 120’ as indicated) at 37°C; 120’ on ice +a.w was used as control for each time point; **E)** Graphical representation of geometric mean of Dara-AF-647 signal in NK cells showing CD38 internalized at 37°C at different time points (15’ to 120’ as indicated); a set of samples incubated 120’ on ice were used as negative control; **F)** Intracellular calcium mobilization in response to Dara (10 μg/ml) was measured by flow cytometry on isolated HD-derived NK cells previously loaded with Fluo-4 AM. Kinetic graph shows an increase of Fluo-4 AM geometric mean as soon as 30 seconds after addition of Dara. A 40% increase was observed at time range 4 of acquisition (103-137 seconds) compared to the baseline time range 1 (0-30 seconds). **G)** Western Blot analysis showing CD38 levels in primary NK cells treated for 6 hrs with Dara (10 µg/ml) or C-IgG, showing LC3II protein downregulation and GLUT4 upregulation; The experiment was performed in at least n=3 independent donors; **H)** Western Blot analysis showing CD38 levels in primary NK cells treated for 16 hours with rapamycin as indicated in the presence of either C-IgG or Dara; **I,J)** Western Blot analysis showing CD38 levels in primary NK cells treated for 16 hours with wortmannin (Wort.) as indicated in the presence of either C-IgG or Dara. The experiment was repeated in biological triplicate, and CD38 relative density levels (CD38/β-actin) were normalized to C-IgG–treated NK cells for each HD donor, the differences reported **(J)**; **K)** Flow cytometry analysis showing that Wort. treatment can partially rescue CD38 surface expression on CD56+CD3(-) NK cells after 6 hrs of Dara treatment.

### Dara-treated CD38+ NK cells induce monocyte activation

Because IFN-γ and GM-CSF release are essential not only for NK cells to directly induce target cell cytolysis but also to indirectly activate the innate immune response, leading to monocyte activation and polarization towards the tumor site (Fig.4A),^27^ we investigated whether patients responding to Dara had an effective increase in monocytic-derived macrophages in their bone marrow. Our data show a significant increase in the frequency of the BM CD14+ population in patients having an effective clinical response, including 1 patient with a partial response (PR), 2 with a very good partial response (VGPR), and 2 with a complete response (CR), compared to that in RRMM and Dara-RRMM (patients that discontinued Dara therapy because of progression) (p<0.001) (Fig.4B). To assess whether NK cells could play a role in this phenomenon, we first tested whether Dara in an ex-vivo model could induce mechanisms of monocyte expansion and polarization. We observed a strong upregulation of costimulatory T cell surface antigens (CD80/CD86) on the surface of the monocytes (CD14+) in PBMCs treated with Dara for 24 hrs (Fig.4C). We then investigated whether the effect of Dara on NK cells may be responsible for inducing markers of monocyte activation (CD80/CD86). CD14+ cells co-cultured with Dara-treated NK cells significantly upregulated CD80/86 (Fig.4D-E) (median average 36%), an effect not observed with C-IgG–treated NK cells (median average 19%, p=0.004, using three different healthy donors). CD80/86 upregulation was not detected in the absence of CD38+NK cells when CD14+ cells were directly exposed to Dara or C-IgG (Fig.4F). However, an increase in the CD80/86-positive population was observed with the addition of 50 ng/mL human recombinant (hr-) IFN-γ for 24 hrs (Fig.4F). Phagocytosis assays show that CD14+ cells incubated with the conditioned media of Dara-treated NK cells for 72hrs display an almost 4-fold increase in CD38+ MM cell death over that from C-IgG–treated NK cells (Fig.4G). As expected, the same phagocytosis effect was observed when isolated CD14+ cells were cultured with hr-IFN-γ at the same time-point (Fig.4G).

**Fig. 4.**
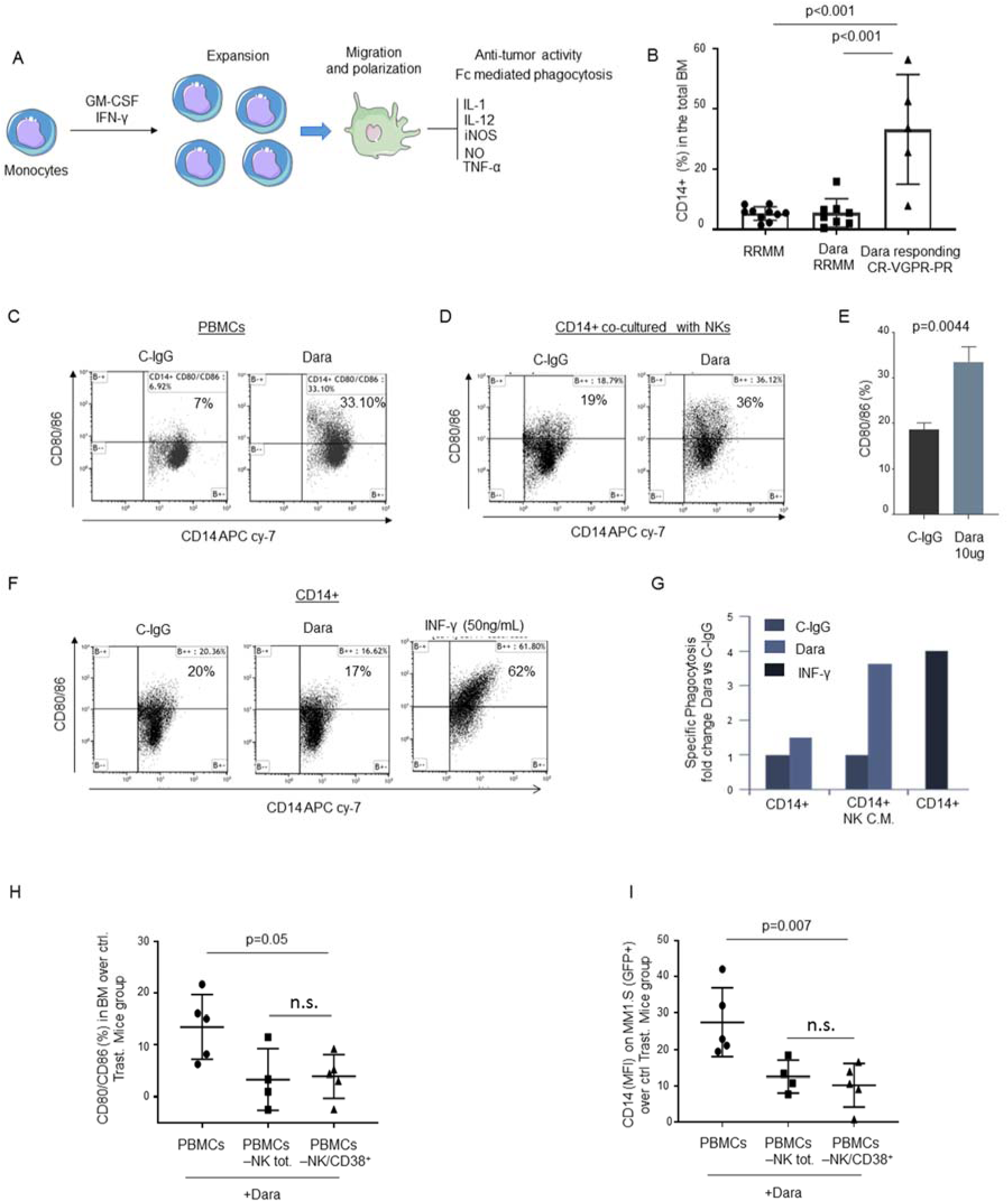
NK cell activation by Dara is essential for monocyte activation and polarization. **A)** Illustration showing monocyte activation and polarization to M1 monocytic-derived macrophages with anti-tumor activity; **B)** Percent of CD14+ cells in the total BM in RRMM, Dara-RRMM, and patients actively responding to Dara; **C)** Representative flow cytometric analysis of CD80/86 in PBMCs isolated from an healthy donor and treated overnight with Dara (10 μg/ml), showing up-regulation of CD80/86 antigens in contrast to that from C-IgG; **D)** Representative flow cytometric analysis of CD80/86 in monocytes isolated from a healthy donor co-cultured with CD38+NK cells (isolated from the same healthy donor) treated overnight with C-IgG or Dara (10 μg/ml); **E)** Bar graph showing CD80/86 upregulation in CD14+ cells co-cultured with Dara-treated NK cells versus C-IgG–treated cells. The experiment was repeated using 3 different donors and represents the average ±SD; p values were calculated using t test, unpaired (tails = 2); **F)** Flow cytometry analysis of CD80/86 in CD14+ cells treated overnight with C-IgG, Dara or INF-γ (50 ng/ml); **G)** Phagocytosis assay was performed on the CD14+ population isolated from a healthy donor, treated for 72hrs with conditioned media (CM) collected from NK cells treated overnight with C-IgG (10 μg/ml) or Dara (10 μg/ml) and compared to the CD14+ isolated population treated for 72hrs with INF-γ (50 ng/ml) at the same timepoint; **H-I)** Dot plot graphs showing the percentage of the CD80/86+/CD14+ population, as assessed by flow cytometry, isolated from the BM of mice injected with total human PBMCs, PBMC depleted of total NK cells, and PBMC depleted of CD38+ NK cells, each fraction pretreated with Dara (**H**); and the increase in GFP+/CD14+ population in the BM of mice injected with total PBMCs versus the other groups (**I**); The hCD14 presence in the BM upon Dara treatment for each treatment group was normalized using the corresponding Trast-treated total PBMCs, PBMC depleted of total NK cells, and PBMC depleted of CD38+ NK cells, respectively. Data reported in **H** and **I** represent the mean ± SD; p values were calculated using multi comparison one-way ANOVA.

The effect of NK cells on monocyte activation upon Dara treatment was also tested in NSG mice engrafted with CD38+ MM cells (MM.1S Gfp/Luc+) (Fig.4H-I). We found a significant upregulation of the CD80/86 marker on the surface of human CD14+ (hCD14+) cells isolated from the BM of the mice treated with Dara-treated PBMCs compared to the BM levels found in mice injected with Dara-treated PBMCs that were depleted of total or CD38+ NK cells (Fig.4H). CD80/86 upregulation in Dara-treated PBMCs was associated with increased recruitment of hCD14+ cells in the mouse BM engrafted with MM.1S Gfp+ cells, as evidenced by an increase of the CD14+/Gfp+ double-positive population, compared to that in mice treated with PBMCs depleted of either total or CD38+ NK cells (Fig.4I). The recruitment of hCD14+ cells and the levels of CD80/86 surface markers observed in mice injected with total PBMCs, CD38+NK cell–depleted PBMCs, and total NK cell–depleted PBMCs and treated with C-IgG were used to normalize BM CD14 engraftment for each of the respective Dara-treated groups.

### MM cells of Dara progressing patients retain targetable surface CD38

Having observed that changes in CD38 surface levels in NK cells are associated with a Dara-induced immune response, we decided to investigate whether changes in CD38 expression on the MM cells could be instead involved in mechanisms of resistance. We collected BM aspirates from 8 relapsed MM patients progressing under Dara-based therapy for ≥4 weeks but prior to starting a subsequent therapy (<16 weeks), and whose material passed our quality control (BM total cellularity >1×10^7^; viability >70%, and a minimum of 1% CD138^+^ MM-PCs among total BM cells). Specifically, we analyzed the BM samples of heavily refractory patients who previously failed at least three lines of therapy but clinically responded (duration of response: 5 months to 24 months) to Dara but then progressed under Dara-based treatment (Dara-RRMM) and compared them with an independent cohort of 8 Dara-naive and 2 Dara-exposed (>1 year prior the analysis) heavily refractory patients (RRMM) that at the time of the analysis were progressing under a different therapeutic regimen. To analyze whether CD38 was still present on the surface of CD138+ MM plasma cells (MM-PCs) from Dara-RRMM patients, we used a non-competitive anti-human CD38 antibody (CD38-Multi-FITC), which binds surface CD38 in the presence of Dara (Sup. Fig.3A,B), versus a CD38 monoclonal antibody (CD38-Mono PE) that shares with Dara the same targeted CD38 epitope (clone IB6) (Sup. Fig.3C,D), as confirmed in cell lines and primary samples (Sup.Fig.3A-D). We did not find significant differences in CD38 surface molecules (levels expressed as mean fluorescence intensity [MFI]) or in percent of CD38+ MM cells with respect to Dara-RRMM and RRMM (Fig.5A-C). Next, we studied whether CD38 on the surface of MM cells from Dara-RRMM patients was still recognizable by Dara. We performed ex-vivo Dara treatment of MM cells for 1 hr, after which no CD38+ signal was detected upon staining with a Dara-competitive fluoresceinated antibody (-PE) (Fig.5D,E, Sup.Fig.3E). In contrast, surface CD38 was detected when the non-competitive antibody was used (Fig.5D,F, Sup.Fig.3F). We were unable to identify any missense mutations in 3 out of 5 of the MM-PCs isolated from Dara-RRMM patients, which were used for ex-vivo Dara binding experiments (data not shown). Moreover, by using the MMRF IA9 released data set, we found no CD38 mutational or mRNA downregulation in the MM cells isolated from 7 patients who were analyzed before and after progressing on single-agent Dara. Immunohistochemistry analysis shows CD38 expression in the CD138+ MM-PCs in the BM biopsies of two independent Dara-RRMM patients in whom marrow was collected before Dara treatment. A second marrow was collected for each patient, 1 patient at an early sign of progression (M spike) before reaching progression by IMWG criteria, and 1 patient at the time of effective progression (Fig.5G,H).

**Fig. 5.**
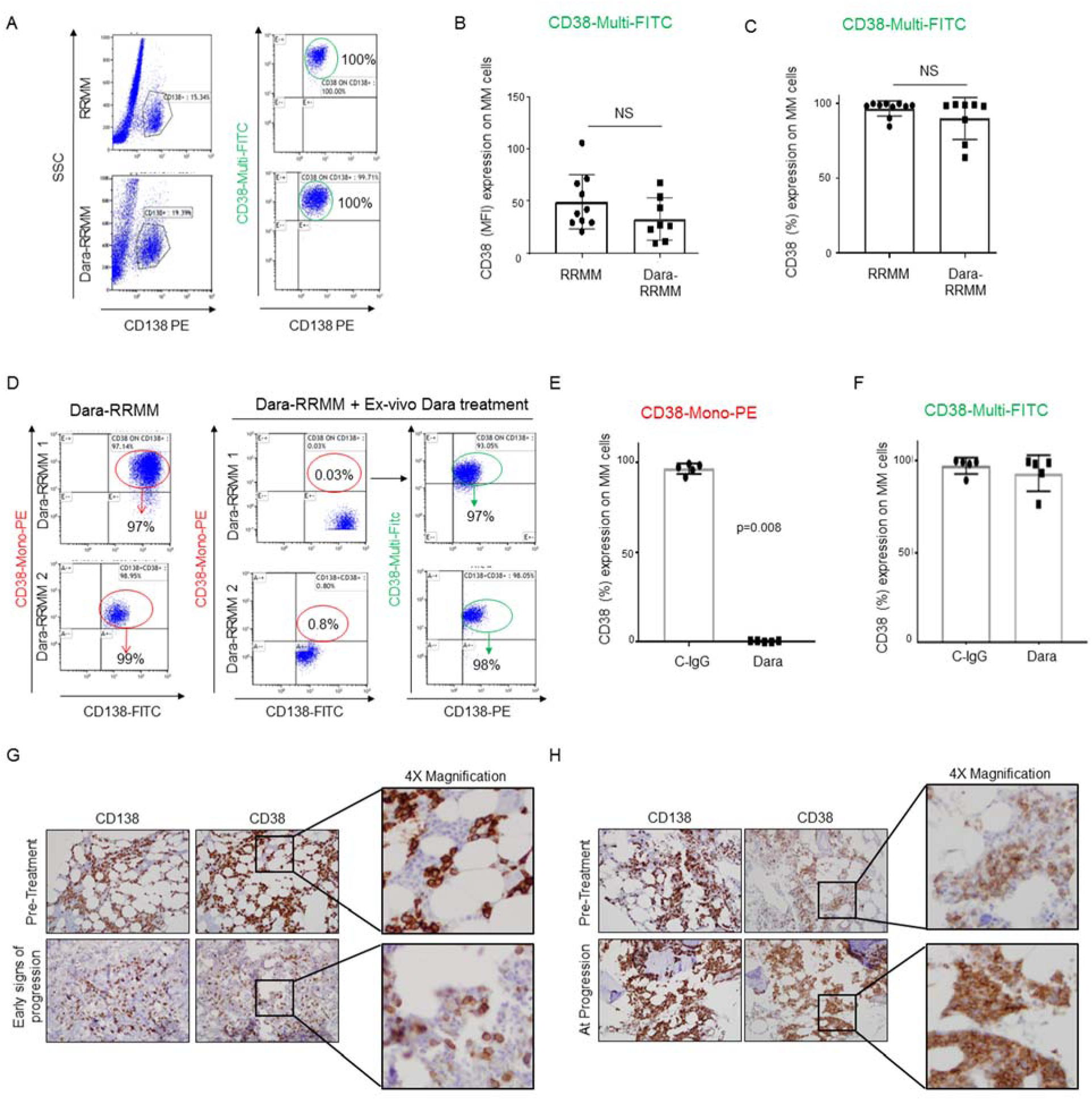
Dara-RRMM patients retain surface CD38 on MM cells. **A)** Representative flow analysis of CD38 expression showing no significant modulation of CD38 expression (%) on CD138+ MM-cells obtained from Dara-RRMM (n=8) compared to RRMM patients (n=10); **B,C)** Bar graphs showing no significant regulation of CD38 expression by MFI **(B)** and % **(C)** among Dara-RRMM and RRMM patients; **D)** Representative flow analysis of CD38 on CD138+ MM cells of two Dara-RRMM patients where BMCs were treated ex vivo with Dara for 1 hr, showing lack of CD38 in MM cells treated with Dara and stained with CD38-Mono PE, and presence of CD38 when the same cells were stained with CD38-Multi PE; **E,F)** Bar graphs showing CD38 surface levels in CD138+ MM cells of 5 Dara-RRMM patients after ex-vivo Dara incubation and stained with CD38-Mono PE **(E)** and CD38-Multi FITC antibodies **(F)**; **G,H)** Immunohistochemical staining for CD138 and CD38 of BM core biopsies or clot sections of 2 MM patients whose BM samples were collected before Dara treatment (pre-Treatment); a second BM biopsy was collected at early signs of progression (G) and at progression (H).

### Dara-RRMM patients retain effector cells with impaired cellular immune function

Although the data from our cross-sectional study (RRMM *versus* Dara-RRMM) show that Dara-RRMM patients maintain targetable CD38 expression on their cancer cells, it could not be excluded that Dara-RRMM patients progressed because of lowered CD38 surface molecules on their cancer cells. To unequivocally assess whether CD38 surface expression on the target cells is essential for Dara-induced cancer cell killing, we tested HL60, a parental leukemia cell line, and HL60 with deletion of CD38 (CD38 CRISPR knockout) (Fig.6A).^28^ Immune cell killing assays show that Dara induced killing in CD38+ HL60 parental (WT) cells, but specific killing was not detected in CD38-negative cells (Fig.6B). We therefore investigated whether Dara-induced MM cell killing could be instead proportional to the MM CD38 surface level expression. We generated five Gfp+/Luc+ MM cell lines having various CD38 surface expression levels (MM.1S, RPMI-8226, KMS11, JJN3 and U266) (Fig.6C). Using the immune-effector cells isolated from the PB of four different healthy donors, we found no significant differences in Dara-induced MM cell killing among these cells regardless of amount of expression of CD38 (MM.1S [75%+], RPMI-8226 [98%], KMS-11 [11%+], JJN3 [9%+]) (Fig.6D). Conversely, specific Dara-induced MM cell killing was not observed in a CD38 surface-negative MM cell line (U266) (Fig.6D). Having observed that high CD38 surface levels on cancer cells are not essential for Dara-induced MM cell killing, we thus assumed that mechanisms of progression could not be related to the complete or even partial loss of CD38 target on MM-cells. Hence, we investigated whether lack of killing of CD38+ MM cells could instead come from changes in the microenvironment of MM patients. A flow cytometry based-killing assay showed that effector cells obtained from Dara-RRMM patients (n=6) incubated with Dara were unable to induce killing of CD38+ MM cells (MM.1S), an effect that was instead observed when effector cells isolated from RRMM patients (n=4) were used (p<0.001) (Fig.6E-F), as was also observed in Dara-responding patients as previously published by Casneuf *et al.*^29^ Mass cytometry analysis (CyTOF) in the PBMCs of Dara-RRMM patients (n=3) that responded to the Dara regiment for more than 18 months (18-22 cycles) and then progressed showed complete absence of the total NK cell population and functional impairment in the CD8+ immune reactive T cell population, as shown by a decrease in HLA-DR and CD127 expression, in contrast to one Dara-responding patient that was treated with 22 cycles of Dara but at the time of the analysis was still in complete remission (Fig.6G) who also displayed an immune signature including cytotoxic T cell expansion and activation that fully resembled the one observed in single agent Dara-responding patients, as recently published (Fig.6H).^16^ Increased expression in the immature monocytic population (CD14+), as assessed by an increase in CD33 and decrease in CD16 expression and an almost 3-fold increase in monocytic myeloid derived suppressor cells (m-MDSCs), was also observed in the Dara-RRMM patient compared to that in the responding patient (Fig. 6H).

**Fig. 6.**
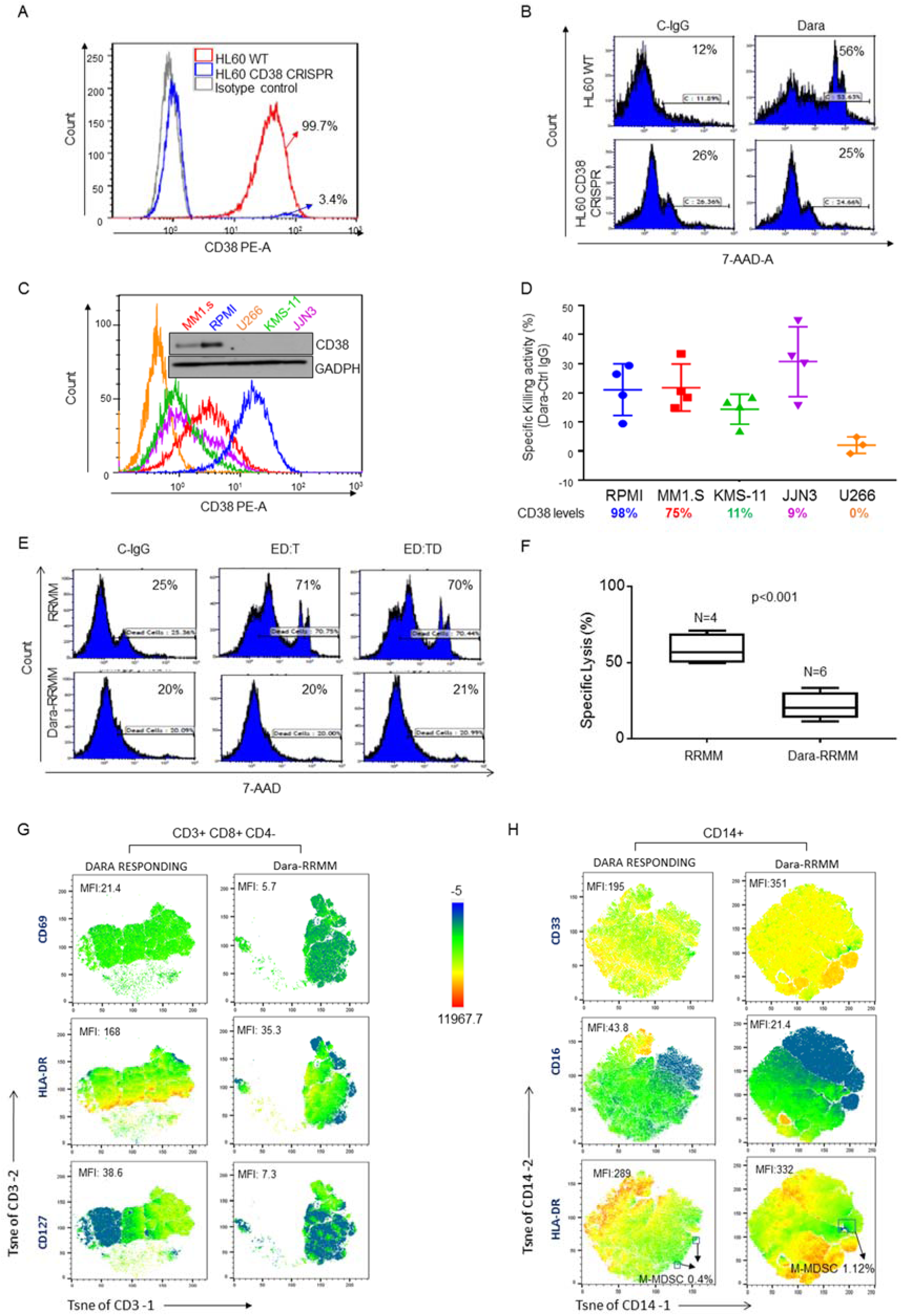
Dara-RRMM patients retain targetable levels of CD38 but impaired immune effector population. **A,B)** Overlay histogram showing CD38 surface expression in parental leukemia cell line HL60 (WT) (99.7%) and HL60 with CD38 deletion by CRISPR (3.4%), compared to the isotype control **A)**, and **B)** representative flow cytometry based-killing assay showing specific killing activity using as target HL60 CD38+ and HL60 CD38-negative cells and as effectors PBMCs from HDs, showing Dara-induced specific killing activity only in the parental cell line (WT); **C)** Overlay histogram showing CD38 surface expression on 5 different Gfp+/Luc MM cell lines (MM.1S, RPMI-8226, KMS11, JJN3 and U266) and Western blot analysis showing CD38 protein level expression. Data are expressed as the mean ± SEM (n=3), normalized compared to control GAPDH; **D)** Representative killing assay of 5 different Gfp+/Luc MM cell lines (MM.1S, RPMI-8226, KMS11, JJN3 and U266) using HD PBMCs as effector cells and treated for 24 hrs with C-IgG or Dara (10 µg/ml or 100 µg/ml), washed, then co-cultured with Gfp+/Luc MM cell lines (target cells) at a ratio of 8:1 for 12 hrs. The graph reports the percentage of dead cells as 7-AAD-positive among target cells of at least n=3 healthy donors repeated in triplicate for each experimental point; **E)** Representative flow analysis of killing assay performed using effector PBMCs of RRMM and Dara-RRMM patients. Effectors were activated for 24hrs with IgG as control, then co-cultured with target cells (MM1.S cells) and treated with Dara. We evaluated percentage of dead cells as 7-AAD positive among target cells (MM1.S GFP+ cells) (ED:T, effectors are treated with Dara; ED:TD, effectors and targets are treated with Dara); **F)** Bar graph showing killing induction (%) in a set of RRMM patients (n=4) and Dara-RRMM patients (n=6) treated with Dara ex vivo. Dara-RRMM patients show impaired killing induction on MM cells (MM1.S) compared to that in RRMM patients (p<0.001); **G)** tSNE heatmap statistic plot obtained by CyTOF analysis showing CD69, HLA-DR and CD127 expression (MFI) in the CD8+ cell subpopulation (gated in total CD3+ cells) of PBMCs isolated from Dara-RRMM and Dara-responding patients; **H)** tSNE heatmap statistic plot obtained by CyTOF analysis showing CD33, CD16 expression (MFI) and frequency of M-MDSC population (gated as CD14+CD11B+CD33+CD15-HLA-DRneg/low in the total CD14+ population) of PBMCs isolated from Dara-RRMM and Dara-responding patients.

## Discussion

Although the role of NK cells in Dara mechanisms of action have been highly debated, here we show that the therapeutic targeting of CD38+NK cells may play a pivotal role in initiating a Th1 immune response,^30, 31^ which can be an essential component in mounting a powerful anti-CD38 immune response, as recently reported by Atanackovic *et al.*^32^ In agreement with Wang et al.,^17^ we demonstrate that Dara induces NK cell degranulation and IFN-γ release, but we show that this effect is restricted to Dara binding to the CD38+ NK cell population, which is indicated to be the only one expressing CD16 on their surface. We additionally observed a considerable increase in CD69 activation marker and GM-CSF production and an increase in Ca^2+^ mobilization in NK cells treated with Dara, which suggests that Dara induces a substantial NK cell activation, a possible explanation as to why we and others observe a significant drop in NK cell frequency in Dara-treated patients.^16^ We also report that in NK cells Dara induces increased autophagy turnover and that CD38 is internalized upon binding. In line with the role of CD38 in immune activation, we observe that the immunosuppressant autophagosome inducer rapamycin downregulates CD38 protein expression in NK cells, whereas the autophagosome inhibitor PI3K inhibitor wortmannin significantly reverts this effect, further supporting the idea that Dara is not selecting a CD38(-) NK population but instead is inducing CD38 degradation to lead to NK cell activation. Interestingly, the levels of CD38 protein expression on NK cells and the magnitude of regulation upon Dara binding appear to be different among different donors (see Fig.3C), an effect that brings us to speculate that differences in the immune system of patients may be critical for the deepness of response to Dara, a hypothesis that will need further validation. In agreement with previous data, which have shown a pivotal role of Dara in inducing macrophage-mediated phagocytosis;^33, 34^ our results demonstrate specifically that direct NK cell activation by Dara induces monocyte activation and polarization, increasing the ability of this subpopulation to kill cancer cells. The macrophage T cell activation markers CD80/CD86, which were upregulated upon Dara treatment via NK cell activation, play a pivotal role not only for macrophage polarization but also as a critical costimulatory molecule in the initial step of T cell activation and survival,^35^ through its binding to the T cell receptor CD28, a hypothesis that is fully consistent with what is observed in Dara-treated patients, as recently reported.^16, 36^ Although we did not perform longitudinal analyses of RRMM and Dara-RRMM patients, we believe the myeloma population analyzed reflects the every-day scenario encountered by clinicians treating MM patients progressing under Dara-based combinatorial therapies. Our data indicate that Dara-RRMM patients retain targetable unmutated CD38 in almost 100% of the MM-PCs, but retain circulating effector cells which are impaired in their killing ability, further supporting the importance of Dara in inducing an anti-MM activity by acting directly through the immune system.

In summary, we propose that Dara induces NK cell activation and degranulation, an effect that can reduce their number but also leads to increased expression of CD80/CD86 T cell costimulatory molecules on monocytes, which induces monocyte polarization in macrophages with anti-tumoral activity and stimulates T cell expansion through CD28 binding, further associated with an immune cascade targeting MM cells (Fig.7). We cannot exclude that Dara can also induce CD38 internalization in the monocyte fraction, with concomitant polarization that may drive a specific anti-CD38 T cell response, a hypothesis that needs further study. In summary, we report that CD38+NK cells can be considered an unexplored therapeutic target which can prime the immune system not only of myeloma patients but also of patients with other forms of cancer. Since anti-CD38 antibodies with toxic payloads and CAR-T cells against CD38 are now under study, the presence of CD38 on MM cells in these patients and a decreased presence in the cancer microenvironment can provide the scientific rationale to use novel CD38-targeted therapeutic interventions in patients progressing on Dara.

**Fig. 7.**
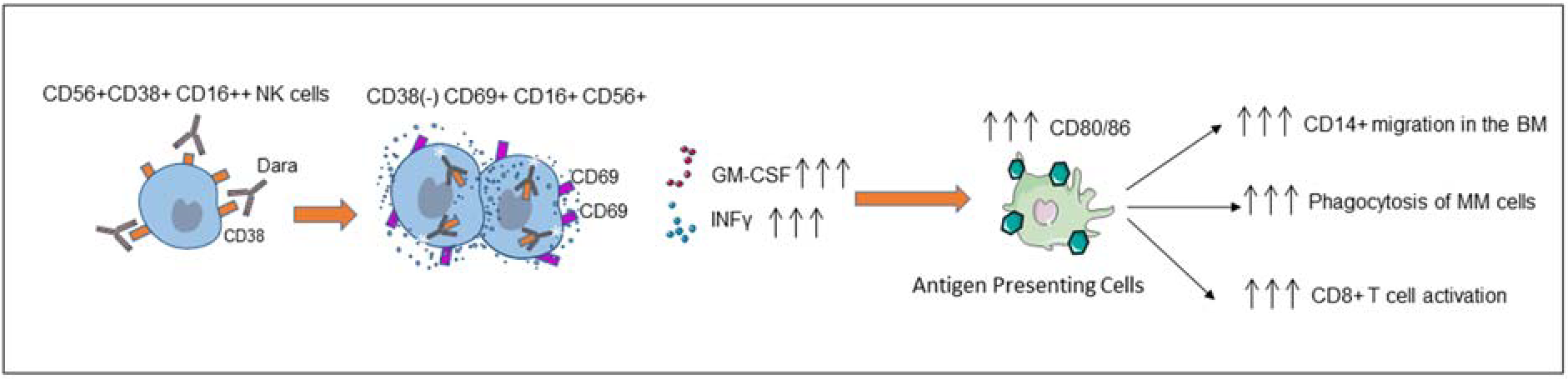
Mechanism of action showing that Dara induces NK cell activation and degranulation and leads to increased expression of CD80/CD86 T cell costimulatory molecules on monocytes, which induces monocyte polarization in macrophages with anti-tumoral activity and stimulates T cell expansion through CD28 binding, further associated with an immune cascade targeting MM cells.

## Materials and Methods

### Flow cytometry

For CD38 expression analysis in MM cell lines and primary samples, cells were washed with PBS 1X and stained for 30 minutes using CD38 PE (IB60) Mouse Anti-Human (Cat.#130-092-260, Miltenyi Biotec)/CD38 FITC Multiepitope (Cat. #CYT-38F2, Cytognos) alone or in combination with CD138 FITC Mouse Anti-Human (Cat. #552723, BD Pharmingen) or CD138 PE (Cat. #561704, BD Pharmingen), CD14-APC Cy7 Mouse Anti-Human (Cat. #557831 BD Pharmingen), CD56 APC (Cat. #362503, Biolegend) CD3 V-450 (Cat. #560351, BD Pharmingen) and CD16 PE cy7 (Cat. #560918, BD Pharmingen) to determine median of fluorescence and percentage of CD38 expression in CD138^+^ MM-PCs, the CD14^+^ monocyte fraction, and the NK CD56+CD3-cell population, respectively, for each patient. CD69 PE (Cat. #310906, Biolegend) and CD107a FITC (Cat. #328606, Biolegend) were used to assess NK cell activation and degranulation. CD80/CD86 PE (Cat. #340294/555658, BD Pharmingen) and CD64 APC (Cat. #561189, BD Pharmingen) were used to determine activation and polarization of CD14+ monocytes. Cells were washed and immediately analyzed on LSRII (Becton Dickinson). Analysis was conducted using Kaluza Software (Beckman Coulter).

### Flow cytometry based-killing assay ex-vivo

Effector cells, freshly prepared PBMCs, NK cells isolated from healthy donors, MM patient-isolated PBMCs and target cells, and GFP+/Luc+MM cells were co-cultured at different ratios (1:1, 4:1, 8:1) and time points. Briefly, effector cells (2×10^6^ cells/well for the 8:1 ratio) were plated at the appropriate concentration then treated, thus activated, with Human-IgG (Cat. #1-001-A) and Dara (Dara-mAb) (Cat. #NCD 57894-502-05) (10 µg/mL or 100 µg/mL) for 24hrs. After incubation, effector cells were washed with PBS 1X solution and then co-cultured with the appropriate ratio of target cells (2.5×10^5^ cells/well for the 8:1 ratio). After 12hrs (4hrs for NK cells) of incubation, the killing induction (%) was assessed by flow cytometry using 7-aminoactinomycin D (7-AAD) (Cat. #51-68981E BD Biosciences).

### Mice experiments

MM.1S GFP+/Luc+ cells were harvested during the logarithmic growth phase and injected intravenously into NOD-SCID mice (0.2 mL/mouse containing 5×10^6^ cells in PBS). Seven days later, mice were monitored daily for tumor development by fluorescence assessment using the In Vivo Imaging System (IVIS). On day 13, after engraftment reached approximately ≥2×10^6^ photons/sec/cm^2^/sr, mice were randomly distributed into the following experimental groups: 1) 1×10^6^ total PBMCs pretreated with trastuzumab (Trast, Cat. #NCD 50242-056-56, 5 mg/kg) (100 µg/mL) (n=6); 2) Total PBMCs pretreated with Dara (n=5); 3) 1×10^6^ PBMCs depleted of total NK + Trast (n=6); 4) PBMCs depleted of total NK + Dara (n=4). Mice were injected once a week for three weeks by intravenous injection. Tumor progression was monitored weekly by fluorescence using IVIS. Injections and treatments for each mouse continued for 3 weeks, which ended when the control groups showed sign of disease.

### Statistics

For *in vitro* experiments, data are reported as mean ± SD of three to four experiments. For ex-vivo Dara incubation studies, blood samples were collected from each HD and were split into the number of treatments under comparison. CD38 expression levels were measured from each treatment group. The Shapiro-Wilk normality test was performed to verify whether the samples were generated from a normal distribution. Appropriate t tests (paired or independent samples; two-sided or one-sided) or nonparametric tests (Mann-Whitney U test, Wilcoxon signed-rank test) were performed to assess significant differences between two (or more) treatments and/or groups. A small p-value (< 0.05) may be considered a statistically significant difference. Animal data were analyzed by ANOVA followed by Tukey’s multiple comparison test for pairwise comparison.

For *in vitro* experiments, data are reported as mean ± SD of three to four experiments. The Student t test (two tailed, unpaired) was used to determine significant differences in experiments within two groups. Animal data were analyzed by analysis of variance (ANOVA). A p value <0.05 was considered statistically significant.

### Study approval

All animal studies were approved by the City of Hope IACUC. Our studies involving human subjects were approved by the IRB of City of Hope and conformed to the tenets of the Declaration of Helsinki. Participants provided informed written consent.

### Primary samples

Total bone marrow aspirates from MM patients and healthy donors (Median age 50-60 years old, male donors), were obtained from City of Hope hematopoietic tissue repository (IRB#16352). Specifically, the cellular fractions of total bone marrow aspirates and PBMCs were isolated using Ficoll-Paque Plus (GE, Healthcare, Life Science) following the manufacturer’s instructions. Fresh BM aspirates from MM patients were routinely depleted from malignant plasma cells using anti-CD138-coated magnetic beads (Cat. #130051301 Miltenyi Biotec) and used for RNA sequencing and functional assays.

### Cell culture, RNA isolation

MM cell lines (MM.1S GFP+/Luc+) were kindly donated from Dr. Irene Ghobrial (Dana-Faber Cancer Institute, Boston, MA). MM cells were cultured in RPMI-1640 medium supplemented with 10% fetal bovine serum (Cat.#019K8420, Sigma), 100 IU/ml penicillin and 100 µg/ml streptomycin. Total cellular RNA was extracted by using TRIZOL reagent (Cat. #15596018 Invitrogen Corporation) and RNA Clean-Up and Concentration Kit (Cat. #43200 Norgen). cDNA synthesis was performed by using the High Capacity cDNA Reverse Transcription Kit (Applied Biosystems, cat# 4368814). Reverse transcription reactions were run using a Mastercycler pro. Quantitative real time-PCR (qRT-PCR) was performed with the TaqMan method (Applied Biosystems), according to the manufacturer’s instructions. The appropriate TaqMan probes for mRNA quantification were purchased from Applied Biosystems, and all reactions were performed in triplicate. The following probes were used: (HS99999905_m1) GAPDH used as endogenous control; (HS01120071_m1) CD38; (HS00989291_m1) INF-γ.

### Immunoblotting

NK cells were washed with phosphate-buffered saline (PBS) and lysate in RIPA buffer (89901, Thermo Scientific) supplemented with protease and phosphatase inhibitors. Lysates were then clarified by spinning at 14,000 rpm at 4^°^C and protein concentration quantified by BCA Protein Assay (23227, Thermo Scientific). Forty micrograms of proteins were denatured in boiling SDS sample buffer and resolved on 4%-20% gradient gels (5671093, Bio-Rad) and transferred to PVDF membrane (1620177, Bio-Rad). After blocking nonspecific binding of antibody with 5% BSA (Fisher BioReagents), blots were probed with one of the following antibodies: anti-CD38 Ab (ab 108403, Abcam), GAPDH (sc-32233, Santa Cruz Biotechnology). Primary antibodies were detected by binding goat anti-rabbit IgG-HRP (NA934, GE Healthcare), goat anti-mouse IgG-HRP (NA931, GE Healthcare), and using an enhanced chemiluminescent visualization system (Cat. #RPN2209 ECL Western Blotting Detection Reagents, GE Healthcare). Primary and secondary antibodies were diluted according to the manufacturer instructions. The bands were quantified by densitometry analyses using Image Lab program (Biorad) and normalized to GAPDH.

### Daratumumab Fab and Fc fragment generation and isolation

The anti-CD38 daratumumab was digested to generate Fab fragments with the proteolytic enzyme IgdE (FabALACTICA Fab kit, Genovis AB, Lund, Sweden). Ten mg of the antibody was digested following the manufacturer’s instructions. The digested antibody was loaded onto a 5 ml CaptureSelect FcXL column (ThermoFisher Scientific, Waltham, MA), equilibrated with 1x PBS and the Fab collected in the flow through. The Fc was eluted using 0.02 M citric acid, 0.05 M NaCl, pH 3.0, into tubes containing 0.1M Tris pH 8.0 (10% v/v). The Fab and Fc were buffer exchanged with PBS and concentrated using Amicon Ultra centrifugal filter (7 kDa mwco, Millipore Sigma, St. Louis, MO).

### Immunohistochemistry analysis

Immunohistochemistry for CD38 (SPC32 clone, 1:100) and CD138 (BA38 clone, 1:1000) was performed on the Ventana Ultra automated immunostainer platform on formalin-fixed, paraffin-embedded tissue sections of bone marrow core biopsies or clot sections prior to and following therapy. Representative fields were photographed at an original magnification of 200x.

### Bioluminescence imaging (BLI) of tumor growth

Non-invasive, whole body imaging for assessment of tumor growth was performed once weekly using the IVIS 100 Imaging system (Xenogen, Alameda, CA). Mice were injected i.p. with 100 μl of the D-Luciferin solution at a final dose of 3 mg/20 g mouse body weight (Biosynth, Cat. No. L-82220) and then gas-anaesthetized with isoflurane (Faulding Pharmaceuticals). Images were acquired for 1–30 sec. images are shown at 60 secs (T0, T1) and 5 secs (T2) from the side angle.

### CD38 internalization into NK cells

To evaluate CD38/anti-CD38 internalization in NK cells isolated from peripheral blood of healthy donors, 1×10^6^ cells were incubated with Dara conjugated with Alexa-Fluor 647 for 15, 60, and 120 min. at 37°C and at 4°C. At the end of the incubation, 5 ml of stripping buffer (100 mM NaCl/100 mM glycine, pH=2.5) was used with incubation on ice for 5 min. NK cells were then spun down (1,000 rpm for 5 min.), washed with PBS (3X) at room temperature, and then analyzed on LSR Fortessa X-20 (Becton Dickinson). Analysis was conducted using Kaluza (Beckman Coulter).

### Dara conjugation

A 20 ml vial of Dara was purchased from the City of Hope pharmacy (400 mg/vial), and 0.3 mg of antibody (2 mg/ml in PBS, 145 µL) was reacted with 0.02 mg of NHS-AlexaFluor647 (2 mg/ml in 50 mM sodium bicarbonate, pH 8.3, 10 µL) for 1 h at room temperature. The reaction was quenched by the addition of 20 µL of 1 M Tris, pH 8.0, and the product was separated from unreacted substrate on a Zeba column (7K MWCO, Thermo Fisher) equilibrated with PBS.

### CyTOF analysis

A total of 1-3×10^6^ PBMCs, derived from Dara responding and Dara-RRMM patients were incubated with a mixture of our custom-made 28 surface cellular marker antibodies. Samples were acquired on a Helios, and data were collected in an FCS file format and normalized using CyTOF Software 6.5.358 for Stand-Alone Processing Workstations (Fluidigm). FCS files are uploaded in FlowJo software for further manual gating and tSNE plot analysis for T cells and monocytes.

## Acknowledgments

We thank Dori Triplet, Evelyn Flores and Debbie Flood for administrative support. Research was in part supported by the National Institute of Health under grant number NIH-2-R01-CA201382 (FP) and in part by the Steven Gordon and Briskin Family Innovation Grant. Research reported in this publication included work performed at the Liquid Tissue Bank, Analytical Cytometry, and Integrative Genomics and Bioinformatics shared resource cores supported by the National Institutes of Health under award number P30CA033572. The content is solely the responsibility of the authors and does not necessarily represent the official views of the National Institutes of Health.

## Author contributions

DV, AD, SJF, PJY, JJK, JS, STR, GM, AK, and FP designed research; DV, AD, MM, EGG, ET, EC, FB, X Wu, SB, TM, ST, CK, MH, SW, MR, DS, RRM, TE performed research; DV, AV, LG, JFS, LL, NC, X Wang analyzed data; DV, AD, JFS and FP wrote the manuscript; all authors reviewed the manuscript.

