## Supplementary Materials for "Daratumumab induces mechanisms of immune activation through CD38+ NK cell targeting"

### Supplementary Figures

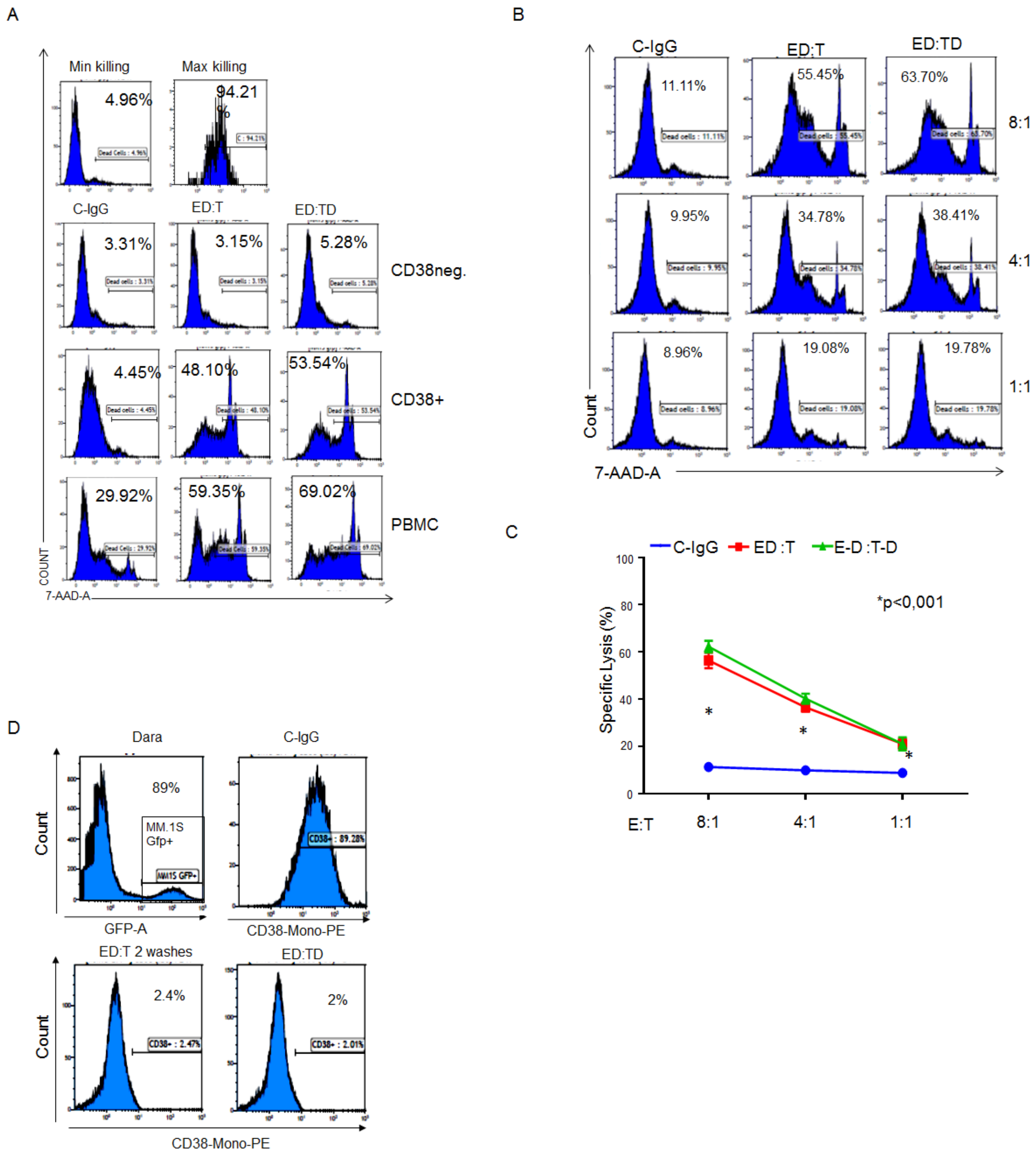

**Fig. S1. CD38+ effector cells. A)** Representative flow cytometry based-killing assay of CD38+ and CD38neg fraction from PBMCs treated overnight with either Dara or C-IgG, washed, and then co-cultured with MM1.S at the ratio of 8:1 for 12h. Analysis in terms of percentage of dead cells as 7-AAD positive among target cells (MM1.S GFP+ cells). Maximum and minimum killing were performed for each set of experiments as internal control for flow cytometry based-killing assay (top dendrograms); **B)** Representative flow cytometry based-killing assay using NK cells (effector cells) isolated from a healthy donor (HD), treated for 24h with C-IgG or Dara, washed, and co-cultured with MM1.S Gfp+/Luc+ (target cells) for 4hr at different E:T ratios, showing equivalent Dara-induced MM cell killing either when the target cells were directly (TD) or indirectly (ED:T) exposed to Dara; **C)** Bar graph reporting the specific lysis (%) of dead cells as 7-AAD positive among target cells (MM1.S GFP+ cells) compared to that from C-IgG. Results are representative of the average of three independent experiments using PBMCs isolated from n=3 healthy donors; **D)** Representative flow cytometry showing complete occupancy of CD38 on the surface of CD38+ GFP+ MM cells (MM.1S) after incubation with PBMCs pre-treated with Dara, washed twice, and incubated with CD38+ MM cells. CD38 occupancy on MM cells was assessed after 1 hour of PBMCs:MM cell co-culture (ED:T) and compared with CD38 occupancy on MM cells treated with PBMCs incubated with C-IgG or MM cells directly exposed to Dara (ED:TD).

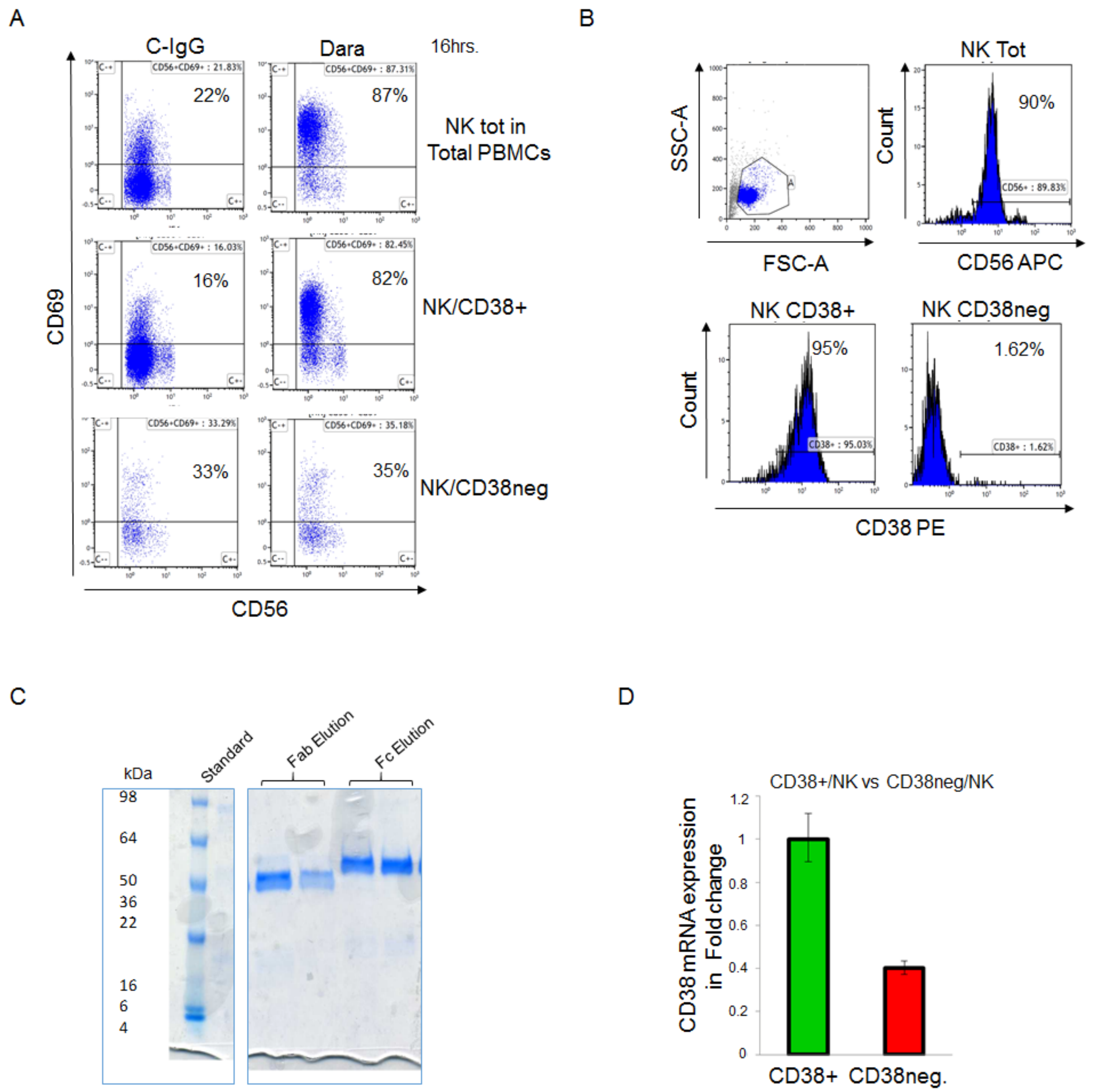


**Fig. S2. Dara induces NK cell activation. A)** Flow cytometry analysis of CD69 expression in the NK cell population (CD56+/CD3-) in total PBMCs, CD38+PBMCs, or CD38-negative PBMC fractions upon overnight treatment with either Dara or C-IgG; **B)** Flow cytometry analysis showing the purity of the fractions: NK total (CD56+/CD3-), CD38+/NK (CD56+/CD3-/CD38+) and CD38-/NK (CD56+/CD3-/CD38-) isolated from the PB of an HD; **C)** SDS-Page gel showing the purity and the size of Dara Fab and Fc fragments after digestion and size exclusion chromatography; **D)** CD38 mRNA baseline expression levels in CD38+/NK cells vs CD38neg/NK cells. CD38 mRNA levels between the two NK cell populations were normalized using GADPH as housekeeping gene.

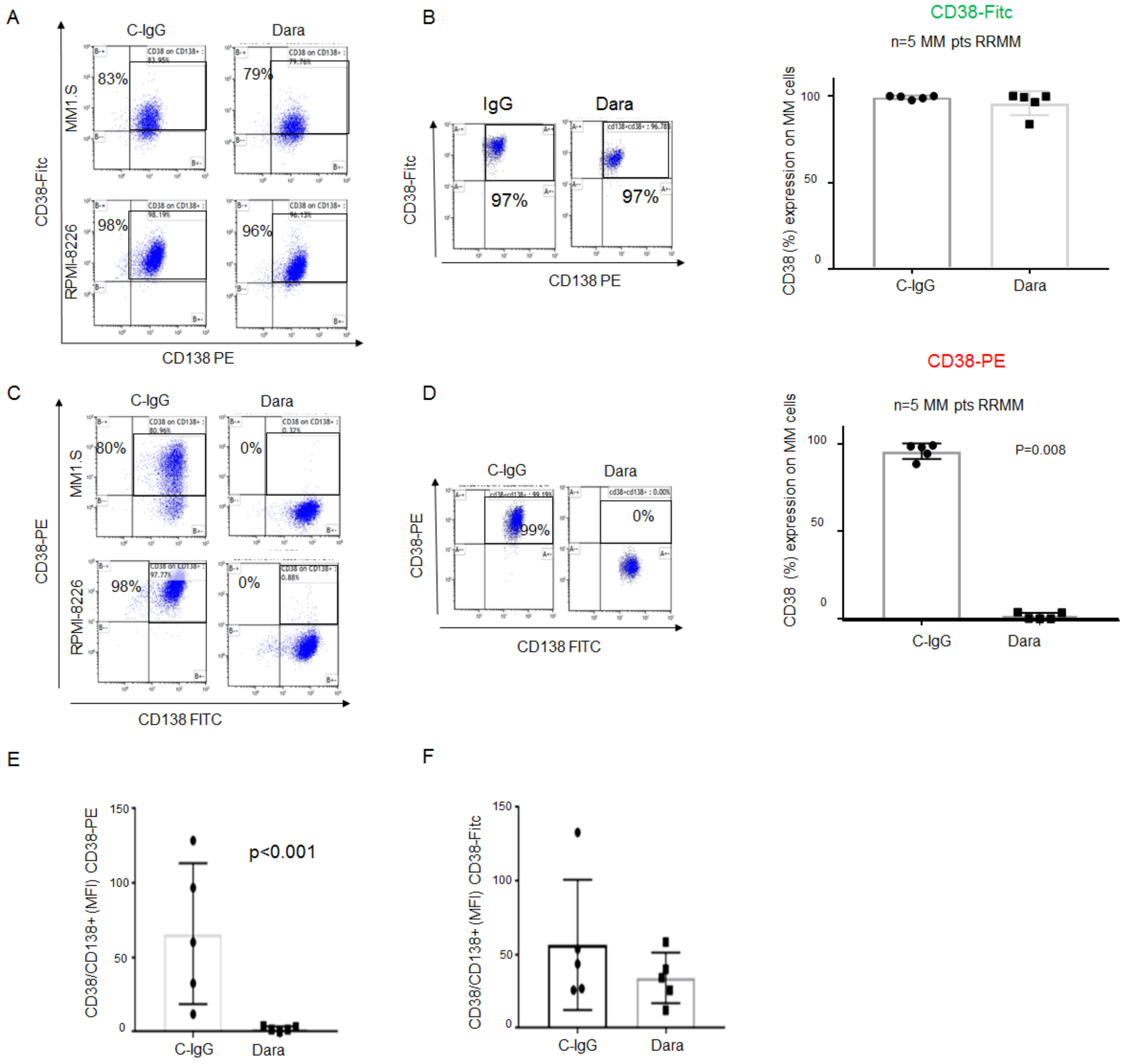


**Fig. S3. CD38 expression in MM cells upon Dara treatment. A)** Representative flow analysis showing no significant modulation of CD38 expression under Dara treatment by using anti-CD38 Multi FITC Ab on MM cell lines MM1.S and RPMI-8226; **B)** Representative flow analysis (left site) of CD38 expression in CD138+ MM cells obtained from a Dara-naïve patient treated ex-vivo for 1 hr with Dara (100 µg/ml) and stained with anti-CD38-Multi Fitc. Bar graph showing comparable CD38 expression between C-IgG and Dara ex vivo treatment detected on MM cells isolated from 5 Dara-naïve pts; **C)**

Representative flow analysis of CD38 showing a significant modulation of CD38 expression in MM cell lines (MM.1S, RPMI-8226) by using anti-CD38-Mono PE in cells treated with Dara versus human control IgG (C-IgG); **D)** Representative flow analysis (left site) of CD38 expression in CD138+ MM cells obtained from a relapsed/refractory multiple myeloma (RRMM) patient treated ex-vivo for 1 hr with Dara (100 µg/ml) and stained with anti-CD38-Mono PE (left). Bar graph (right) showing dissimilar CD38 expression between C-IgG and Dara ex vivo treatment detected on MM cells isolated from 5 Dara-naïve patients; **E-F)** Bar graph showing CD38 surface expression on MM cells isolated from 5 Dara-RRMM patients, after ex-vivo incubation with Dara or C-IgG and stained with anti-CD38-Mono PE (E) or Multi-FITC (F) and reported in MFI.
